# RNA is required for the maintenance of multiple cytoplasmic and nuclear membrane-less organelles

**DOI:** 10.1101/2021.08.16.456519

**Authors:** Carolyn J. Decker, James M. Burke, Patrick K. Mulvaney, Roy Parker

## Abstract

Numerous membrane-less organelles composed of a combination of RNA and proteins, referred to as RNP granules, are observed in the nucleus and cytoplasm of eukaryotic cells, including stress granules, processing bodies, Cajal bodies, and nuclear speckles. An unresolved issue is how frequently RNA molecules are required for the maintenance of RNP granules in either the nucleus or cytosol. To address this issue, we degraded intracellular RNA in either the cytosol or the nucleus by the activation of RNase L and examined the impact of RNA loss on several RNP granules. Strikingly, we find the majority of RNP granules, including stress granules, processing bodies, Cajal bodies, nuclear speckles and the nucleolus are altered by the degradation of their RNA components. In contrast, super-enhancer complexes and TIS granules were largely unaffected by widespread intracellular RNA degradation. This highlights a critical and widespread role of RNA in the organization of many, but not all, RNP granules.

**SUMMARY:** Decker et al. examine the role of RNA in maintaining membrane-less organelles composed of RNA and protein by activating RNase L in the cytoplasm or nucleus. Degradation of RNA alters the majority of membrane-less organelles indicating that RNA plays a widespread role in their organization.

## INTRODUCTION

Eukaryotic cells contain many different compartments in the cytosol and nucleus that are not bound by membranes (Spector, 2006; Gomes and Shorter, 2019). Many of these membrane-less organelles are large assemblies of RNA and protein, referred to as ribonucleoprotein (RNP) granules. RNP granules found in the cytoplasm include processing bodies or P-bodies (PBs) (Jain and Parker, 2013), stress granules (SGs) (Anderson and Kedersha, 2006), U-bodies (Liu and Gall, 2007), IMP granules (Jønson et al. 2007), FMRP granules (Mazroui et al., 2002), TIS granules (Ma et al., 2018), neuronal granules (Keibler and Bassell, 2006) and germinal granules (Voronina et al., 2011). Examples of nuclear RNP granules include the nucleolus, nuclear speckles, paraspeckles, Cajal bodies and nuclear stress granules (Biamonti, 2004; Mao et al., 2011). Understanding how these non-membrane-bound compartments or organelles assemble and maintain their integrity will provide fundamental insight on the organization of eukaryotic cells.

The role of RNA in organizing these compartments in the cell is an important question. There is evidence that RNA is required for the formation of some of these organelles or chromatin compartments. In the case of paraspeckles a long non-coding RNA (lncRNA), NEAT1_2, plays a core architectural role in assembly (Clemson et al., 2009; Sasaki et al., 2009). At the end of mitosis, nucleoli form around the sites of transcription of precursor rRNA and nucleolar assembly also requires 45S rRNA produced prior to mitosis (Hernandez-Verdun 2011). Formation of SGs and PBs in the cytoplasm requires non-translating mRNAs (Kedersha et al., 1999; Teixeira et al., 2005; Buchan et al., 2008; Buchan and Parker, 2009). Whether RNA plays an essential role in the formation of other RNP bodies is less clear. For example, a long non-coding RNA called MALAT1 is enriched in nuclear speckles, but it is not required for nuclear speckles to form (Hutchinson et al., 2007). However, poly(A)+ RNA is present in nuclear speckles under conditions where MALAT1 RNA localization to speckles is defective (Miyagawa et al., 2012) so whether other RNA species are involved in organizing nuclear speckles remains to be determined. Whether RNA is required for the stable maintenance of membrane-less organelles once they are formed is also an unresolved issue.

One way that RNAs could promote RNP body assembly is by being a scaffold for RNA binding proteins which then bind to additional proteins either through homotypic or heterotypic protein-protein interactions and/or additional RNAs to form higher order assemblies (Fox et al., 2018). In addition, intermolecular RNA-RNA interactions can directly promote the assembly of RNP granules (Van Treeck et al., 2018). Thus, RNP granule formation can be thought of as the summation of RNA-RNA, RNA-protein and protein-protein interactions (Van Treeck and Parker, 2018). The contribution of these three classes of interactions to the formation and stable maintenance of RNP assemblies may differ between different RNP granules or for the same type of RNP granule under different conditions.

We recently reported that SGs and PBs are differentially sensitive to the loss of RNA in the cytoplasm (Burke et al. 2020) suggesting that the role of RNA-RNA or RNA-protein interactions on their structural integrity differs. The antiviral endoribonuclease, ribonuclease L (RNase L), is activated in response to dsRNA which leads to widespread degradation of mRNAs in the cytoplasm (Burke et al. 2019, Rath et al. 2019). RNase L activity limits the formation of SGs and can disassemble preformed SGs (Burke et al. 2020). In contrast, RNase L activity does not dramatically affect the size of PBs (Burke et al. 2020). The differential response of SGs and PBs to loss of RNA in the cytoplasm suggests that SGs are strongly dependent on RNA-RNA and/or RNA-protein interactions whereas PBs maybe more dependent on protein-protein interactions. Alternatively, RNAs within PBs may be stably associated and protected from RNase L in comparison to RNA associated with SGs.

To better understand the role of RNA in the organization of cellular compartments we have examined whether a variety of RNP granules in the nucleus and the cytoplasm are dependent on the presence of RNA for their integrity. We quantified the effect of RNase L activation in the cytoplasm on the number and volume of PBs and on the integrity of TIS granules. We also targeted RNase L to the nucleus to induce the degradation of nuclear RNA and determine if nuclear bodies maintained their structure or were disassembled. We find that RNAs within nuclear RNP granules are susceptible to RNase L degradation and that granules disassemble, or form assemblies with altered morphology, in response to the loss of their resident RNAs. Taken together, our findings reveal that RNA plays an important role in organizing multiple cellular compartments.

## RESULTS AND DISCUSSION

### P-body number but not size is reduced by RNase L activity

RNase L activity has been reported to inhibit the assembly of SGs but have less effect on PBs (Burke et al., 2019; Burke et al., 2020). Similarly, we observed that SGs assembled in RNase L knockout (RL-KO) A549 cells in response to poly(I:C), a viral dsRNA mimic that activates the RNase L pathway, but their assembly was prevented in wild-type A549 cells with active RNase L, as judged by depletion of a cytoplasmic RNA (GAPDH RNA) (Figure 1A and B). In WT cells, only small G3BP foci, referred to as RNase L-dependent bodies (RLBs) (Burke et al. 2019), assembled (Figure 1A and B).

**Figure 1.**
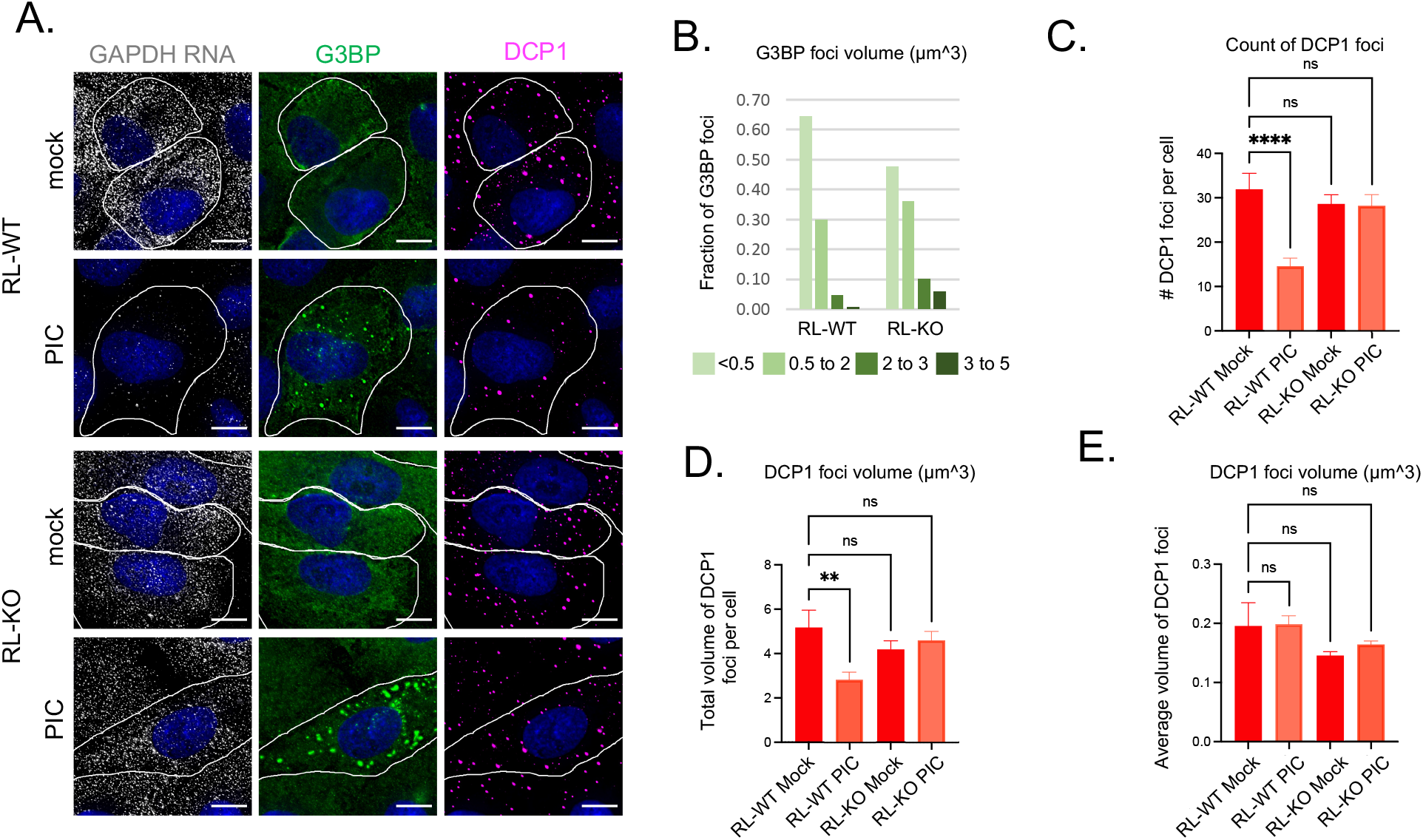
The number of PBs is reduced in cells with active RNase L. A. GAPDH smFISH and IF analysis using anti-G3BP antibody (G3BP) and anti-Dcp1b antibody (DCP1) in A549 cells (RL-WT) or A549 RL-KO cells either mock transfected (mock) or transfected with poly(I:C) (PIC) for 5 hours. Scale bar 10 microns. B. Graph of the fraction of G3BP foci with different volumes in PIC treated RL-WT and RL-KO cells. C. Number of DCP1 foci in mock and PIC treated cells. D. Total volume of DCP1 foci in mock and PIC treated cells. E. Average volume of Dcp1 foci in mock and PIC treated cells. C-E. Mean and SEM of at least 10 cells of each cell type and condition. One way Anova with Dunnett’s multiple comparison. ns non-significant, ** P value ≤0.01, **** P value ≤0.0001.

We observed a clear, but different, effect of RNase L activation on P-bodies. In poly(I:C) treated cells that activated RNase L, there was a 2-fold reduction in PB number and total PB volume compared to RNase L knock-out cells (Figure 1C and D). However, the average size of PBs did not change dramatically in response to poly(I:C) in the RL-WT cells (Figure 1E) (Burke et al., 2020). Targeting RNase L to PBs by fusing it to a PB protein, Dcp1 (Ingelfinger et al. 2002), did not further decrease the number or size of PBs (Figure S1) indicating that the relatively small effect of activating RNase L on PBs is not due to RNase L being unable to access PBs. The reduction in PBs with RNase L activation is consistent with RNA promoting their assembly. However, the unchanged size distribution of PBs implies that features in addition to RNA abundance specify a preferential PB assembly size.

### Degradation of cytoplasmic RNA by RNase L does not disrupt TIS granules

Recent results have identified a new type of RNP granule, referred to as TIS granules, that contains an RNA binding protein, TIS11B, and mRNAs that encode membrane proteins and have 3’UTRs with multiple AU-rich elements (Ma et al. 2018). They form a mesh-like network that is associated with the endoplasmic reticulum (ER) (Ma et al. 2018), which has been proposed to be generated by extensive intermolecular RNA-RNA interactions between mRNAs in the granules (Ma et al., 2021). To determine if the integrity of TIS granules was dependent on resident RNAs we examined the distribution of TIS11B and an ER marker, protein disulfide isomerase (PDI), after activation of RNase L with poly(I:C). Activation of RNase L was detected by the presence of RLBs using oligo(dT) FISH (Burke et al. 2020).

RNase L activation did not dramatically effect TIS granules, as assessed by the subcellular location of endogenous TIS11B detected with the antibody used by Ma et al. 2018. Specifically, in mock treated cells, TIS11B formed irregular shaped assemblies that overlapped but did not co-localize with the ER, although TIS11B was depleted in regions of the ER where the PDI signal was highest (Figure 2A). After poly(I:C) treatment, PDI was less concentrated in the perinuclear region of cells suggesting that exposure to dsRNA may lead to a down regulation of the rough ER (Figure 2A), which could result from general translation repression due to the activation of protein kinase R or degradation of bulk cytoplasmic mRNA (Burke et al., 2019). The distribution of TIS11B did not change markedly in cells with active RNase L with TIS11B remaining in irregular-shaped assemblies (Figure 2A and B). In cells with active RNase L, there was no change in the number of TIS granules (Figure 2C) or their overall volume (Figure 2D).

**Figure 2.**
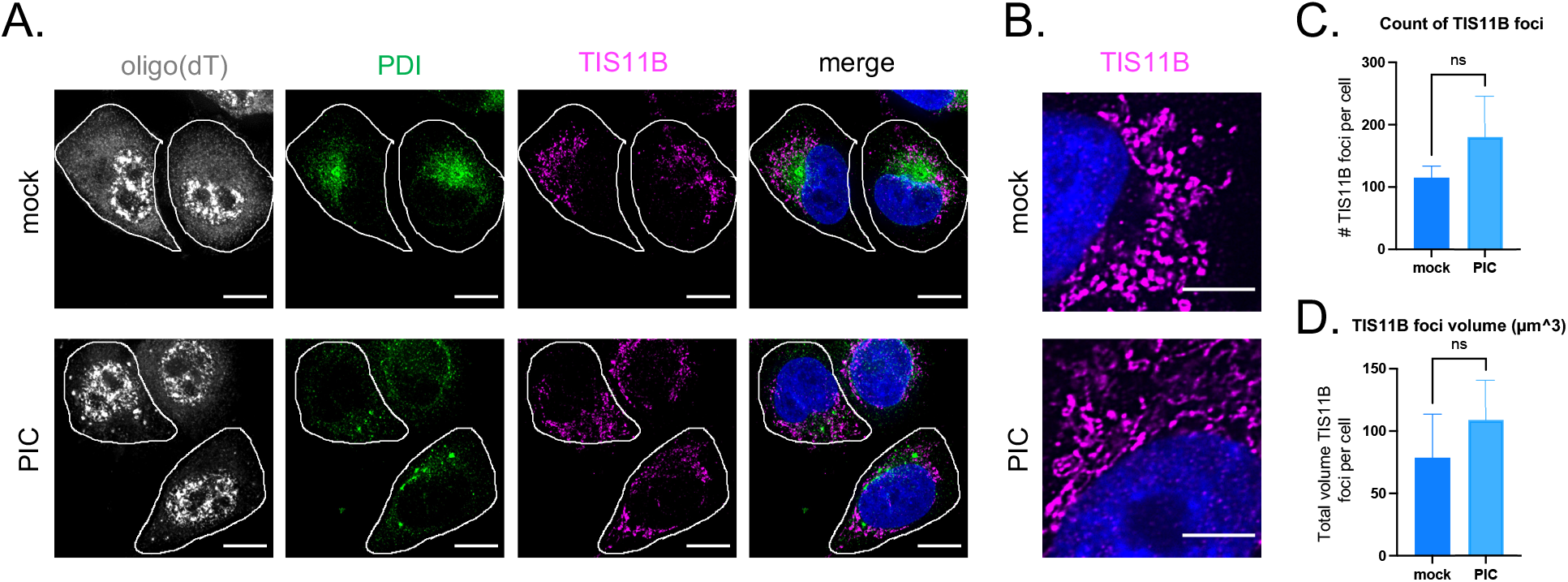
TIS granules are not markedly affected in cells with active RNase L. A. IF analysis using anti-TIS11B antibody or anti-PDI (protein disulfide isomerase) and oligo(dT) FISH to detect poly(A) RNA in mock or PIC treated A549 cells (RL-WT). Scale bar 10 microns. B. Super-resolution images of TIS11 granules in mock or PIC treated RL-WT cells. C. Number of TIS11B foci in mock and PIC treated cells. D. Total volume of TIS11B foci in mock and PIC treated cells. C-E. Mean and SEM of 3 independent experiments with at least 10 cells of each cell type and condition per experiment. Mann-Whitney test, ns non-significant.

The persistence of TIS11B localization after RNase L activation could be because TIS granules do not depend on RNA to maintain their integrity. Alternatively, mRNAs within TIS granules could be resistant to RNase L cleavage. Specific mRNAs that accumulate in TIS granules, at least when overexpressed, include CD47, BCL2, CD274, ELAVL1, CCND1 and FUS (Ma et al., 2018). These mRNAs are present at very low levels (Table S1; Khong et al., 2017), which makes the analysis of their association with TIS granules after RNase L activation impractical without over-expression. Therefore, to examine if these mRNAs are degraded by RNase L we compared their levels by RNA-Seq following RNase L activation (Burke et al., 2019). The levels of the majority of these mRNAs decreased 1.5-3 fold with RNase L activation (Table S1) documenting that these TIS granule-localized mRNAs are sensitive to RNase L degradation.

Another possible explanation for the persistence of TIS11 granules after RNase L activation is the induction of specific mRNAs during the dsRNA response that at least partially escape RNase L degradation (Burke et al., 2019). Many of these induced mRNAs contain AU-rich elements (Burke et al., 2019) and thus may be localized to TIS granules. To test this possibility, we treated cells with Actinomycin D (ActD) to inhibit the production of new transcripts while activating RNase L with poly(I:C) and then monitored TIS granules by immunofluorescence. Inhibition of new transcription did not significantly change the number or overall volume of TIS granules in cells with active RNase L (Figure S2). This result indicates that the maintenance of TIS granules in cells where cytoplasmic RNA is degraded by RNase L is not dependent on the induction of new mRNAs. Taken together, these results are consistent with a model wherein TIS granules do not require RNA to maintain their structure. However, we cannot rule out the possibility that there are one or more specific RNAs required for TIS granule preservation that are resistant to RNase L, and sufficiently stable such that their levels are unaffected during ActD treatment.

### Method to degrade nuclear RNA in an inducible manner

The differential effect of loss of RNA on RNP granules in the cytoplasm led us to test whether RNP granules in the nucleus require RNA to be maintained. We reasoned that we could induce the degradation of nuclear RNA by targeting RNase L to the nucleus and activating its activity through the dsRNA innate immune response.

We fused the c-myc nuclear localization signal (NLS) to RNase L, introduced the modified wild-type RNase L (NLS-RL-WT) and RNase L -R667A catalytic mutant (NLS-RL-CM) into A549 cells using a lentiviral vector, and determined if the NLS was sufficient to target RNase L to the nucleus. By biochemical fractionation, we observed that the NLS-RL-WT and NLS-RL-CM proteins were detected in both the nuclear and cytosolic fractions, while the endogenous RNase L is only in the cytosol (Figure 3A). This demonstrates that the NLS was sufficient to target a portion of RNase L to the nucleus. The exogenous RNase L proteins were overexpressed ~15-25 fold compared to the endogenous protein, which should allow it to robustly degrade both nuclear and cytoplasmic RNAs.

**Figure 3.**
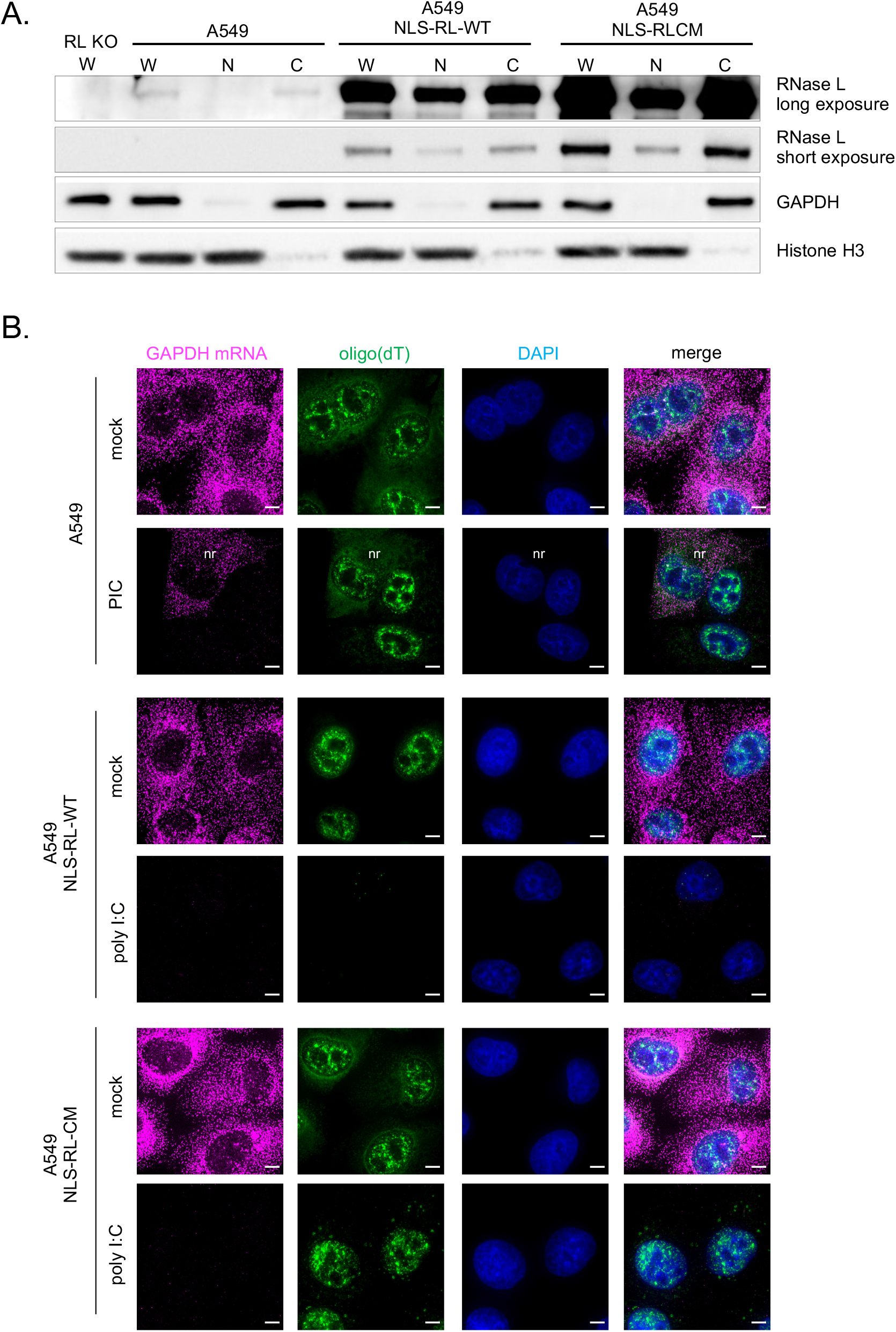
Nuclear-localized RNase L degrades nuclear RNA in response to poly(I:C). A. Western analysis of nuclear (N) or cytoplasmic (C) fractions from whole cell lysates (W) from A549 cells, RL-KO A549 cells, or A549 cells with either RNase L (NLS-RL) or RNase L-R667A (NLS-RL-CM) fused to a nuclear localization signal sequence. Short and long exposure with anti-RL antibody. GAPDH protein used as cytoplasmic marker. Histone H3 protein used as nuclear marker. B. FISH with probes to GAPDH mRNA or oligo(dT) to detect poly(A)+ RNA in A549 cells without or with either NLS-RL or NLS-RL-CM either mock transfected or transfected with poly(I:C) for 4 hours. Scale bar 5 micron. nr indicates cells non-responsive to PIC.

By examining RNAs by FISH, we demonstrated that the localization of RNase L to the nucleus results in RNA degradation in the nucleus. Specifically, nuclear poly(A) signal was strongly lost in cells expressing NLS-RL-WT (Figure 3B) in response to poly(I:C) treatment. The degradation of nuclear poly(A)+ RNAs was dependent on the catalytic activity of RNase L since cells expressing NLS-RL-CM did not degrade nuclear RNA (Figure 3B). Consistent with the tagged RNase L being in both compartments, and the presence of the endogenous RNase L in the cytosol, cytosolic RNA degradation as assessed by loss of the GAPDH mRNA occurred following treatment with poly(I:C) in cells expressing either form of the NLS-RL fusion proteins (Figure 3B). Thus, this system allows the degradation of nuclear RNA in living cells in an inducible fashion. We then used this approach to determine if nuclear RNP granules require RNA to maintain their structural integrity.

### Nucleolar structure is dependent on the presence of RNA

The nucleolus is a large structure in the nucleus where transcription and processing of ribosomal RNAs and their assembly into ribosomal subunits occurs. It is organized into distinct subdomains, a fibrillar center (FC) containing rDNA, the dense fibrillar component (DFC) containing the precursor rRNA and snoRNAs involved in initial steps in rRNA processing and the granular component (GC) where further processing, modification and assembly of ribosomal subunits is thought to occur (Lam et al. 2005). To monitor if nucleolar-associated RNA was accessible to degradation by RNase L, we used a probe for the small nucleolar RNA, snoRD3A, a C/D box snoRNA which localizes to the DFC and GC (Gerbi and Brovjagin 1997). In the mock transfected cells containing NLS-RL-WT, snoRD3A signal was in large nuclear structures (Figure 4A). snoRD3A signal was depleted in cells with activated RNase L in the nucleus, which we identified by loss of nuclear oligo(dT) signal (Figure 4A).

**Figure 4.**
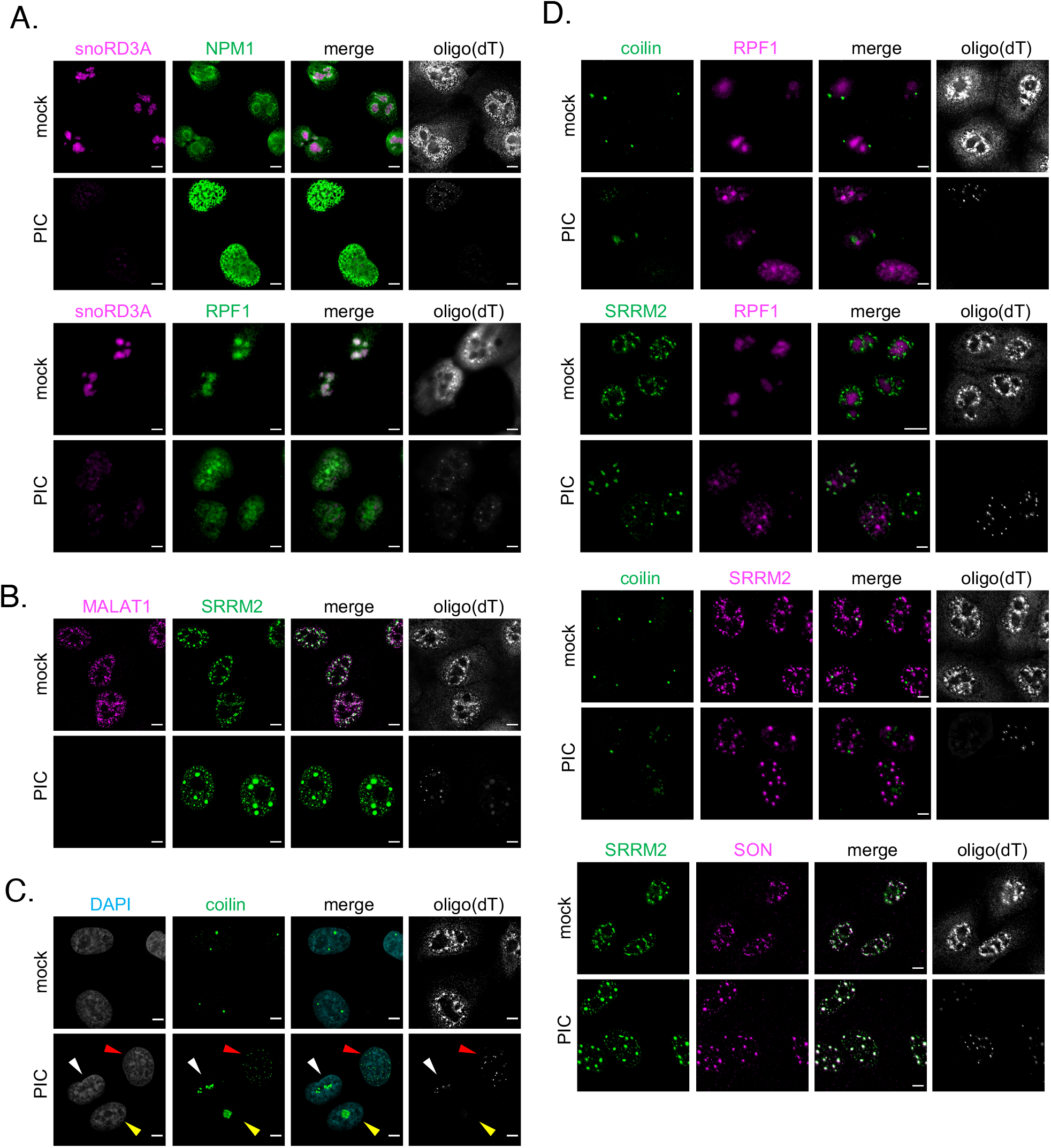
The structure of nucleoli, nuclear speckles and Cajal bodies is dependent on nuclear RNA. Analysis of nuclear RNA granules in A549 cells with RNase L targeted to the nucleus (NLS-RL-WT) either mock transfected or treated with poly(I:C) (PIC) for 4 to 5 hours. Scale bar 5 microns. A. FISH analysis to detect poly(A)+ RNA and nucleolar-localized snoRD3A RNA and IF analysis to detect nucleolar proteins. Top panel: nucleophosmin (NPM1). Bottom panel: ribosome processing factor 1 (RPF1). B. FISH analysis to detect poly(A)+ RNA and nuclear speckle-localized MALAT1 RNA and IF analysis with anti-sc35 antibody to detect nuclear speckle protein SRRM2. C. FISH analysis to detect poly(A)+ RNA and IF analysis to detect Cajal body protein coilin. D. IF analysis of nucleolar protein RPF1, Cajal body protein coilin and nuclear speckle proteins SRRM2 and SON.

Examination of NPM1 (nucleophosmin 1), which is enriched in the GC of nucleoli (Spector et al. 1984), demonstrated that the GC component of nucleoli is lost when nuclear RNA is degraded. In the mock transfected cells, NPM1 was found in the nucleoplasm and concentrated in a ring surrounding the snoRD3A RNA signal (Figure 4A). In cells in which nuclear RNA was degraded, based on depletion of oligo(dT) and snoRD3A RNA signal, NPM1 was dispersed in the nucleoplasm (Figure 4A), suggesting that the granular component of the nucleolus had been disrupted.

We also examined how RPF1 (ribosome production factor 1) was affected by nuclear RNA degradation. RPF1 is localized to nucleoli where it binds to pre-60S ribosomal subunits (Wehner and Baserga, 2002; Kater et al. 2017). RPF1 co-localized with the snoRD3A RNA in the mock transfected cells (Figure 4A). In the majority of cells (70%) with nuclear RNA degradation, RPF1 was dispersed in the nucleoplasm (Figure 4A) consistent with nucleolar structure requiring RNA. We did observe that in 30% of the cells in which the snoRD3A RNA was depleted, RPF1 was concentrated in one or a few large foci (Figure 4A) as well as being found in the nucleoplasm. These residual RPF1 assemblies could be arising in nuclei where RNA degradation is not as robust (see below). No change in RPF1 localization was observed in response to poly(I:C) in cells expressing the nuclear-targeted catalytic mutant form of RNase L (Figure S3A) consistent with the changes in nucleolar structure being dependent on the degradation of nucleolar RNA. Taken together these findings suggest that RNA is required to maintain the overall structure of nucleoli. The requirement for RNA for nucleolar structure is consistent with previous work showing transcription of rRNA is required for nucleolar formation (Benavente, 1991).

### The size and shape of nuclear speckles changes following nuclear RNA degradation

Mammalian cells contain 10-50 irregular shaped nuclear bodies referred to as nuclear speckles that are found in interchromatin spaces within the nucleus (Spector and Lamond 2011). Nuclear speckles contain poly(A)+ RNA and factors involved in mRNA metabolism, including snRNPs, splicing factors, RNA export and quality control factors, subunits of RNA polymerase II, transcription factors and translation initiation factors (Hall et al. 2006, Spector and Lamond 2011). We monitored MALAT1 RNA, a lncRNA found in nuclear speckles (Hutchinson et al. 2007), to determine if RNA in speckles is susceptible to degradation by RNase L. The SC35 antibody, which recognizes the SRRM2 protein (Ilik et al., 2020), was used to monitor the effect of nuclear RNA depletion on nuclear speckles.

We observed that degradation of nuclear RNA, including MALAT1, led to a dramatic rearrangement of nuclear speckles. Specifically, in mock treated cells, we observed polyA+ RNA and SRRM2 protein in discrete nuclear speckles that were often associated with local concentrations of the MALAT1 (Figure 4B). In response to poly(I:C) treatment, MALAT1 and nuclear polyA+ RNA was depleted in cells expressing nuclear-localized RNase L (Figure 4B). Strikingly, the degradation of nuclear RNA corresponded with the loss of nuclear speckles and re-organization of SRRM2 into a few large discrete nuclear assemblies. No change in SRRM2 localization was observed in response to poly(I:C) in cells expressing the nuclear-targeted catalytic mutant RNase L (Figure S3B).

The change in size and morphology of nuclear speckles in response to nuclear RNA degradation suggests that RNA normally limits the assembly of SRRM2 and potentially other speckle components into larger structures. Since MALAT1 is not required for nuclear speckles, and depletion of MALAT1 from speckles does not result in large round bodies (Miyagawa et al., 2012), we infer that RNAs other than MALAT1 determine the morphology of nuclear speckles. Consistent with RNA being required for the morphology of nuclear speckles, nuclear speckle components rearrange into similar large spheroid bodies when transcription is inhibited with ActD (Shav-Tal et al., 2005; Sasaki et al., 2009). This is consistent with the hypothesis that nascent transcripts limit the assembly of speckles.

### Coilin, a Cajal body protein, re-localizes in response to nuclear RNA degradation

Cajal bodies are sub-compartments in the nucleus involved in the assembly and RNA modification of small nuclear ribonucleoprotein complexes required for pre-mRNA splicing, ribosome biogenesis, histone mRNA processing and telomere synthesis (<u>Staněk</u>, 2016; Machyna et al., 2013). In addition to these substrate small nuclear ribonucleoprotein complexes, Cajal bodies contain many protein factors as well as small Cajal body specific RNAs (scaRNAs) (Machyna et al. 2013). To determine if RNA is important for maintaining Cajal body assembly, we examined the organization of Cajal bodies in response to the degradation of nuclear RNA.

We observed that nuclear RNA degradation led to dramatic rearrangement of the Cajal body protein coilin. In mock treated cells, coilin was localized to a few foci per nucleus (Figure 3C). In contrast, in cells in which nuclear RNA was degraded, as assessed by loss of oligo(dT) signal, coilin was found in a variety of different types of assemblies (Figure 4C). In 13% of nuclei, coilin formed one or two large irregular spherical shaped foci (Figure 4C, yellow arrow). Coilin was in irregular shaped foci of varying sizes and number (Figure 4C, white arrow) in 29% of the nuclei. In 23% of the nuclei in which RNA had been degraded, coilin was localized to many small foci that may represent sub-assemblies of Cajal bodies (Figure 4C, red arrow). In 8% of the nuclei without oligo(dT) signal, coilin was dispersed in the nucleoplasm. In the remaining nuclei in which poly(A)+ RNA had been degraded, coilin was found in foci that resembled Cajal bodies in size and number. These foci could be Cajal bodies in which the resident RNAs have not been degraded. The distribution of coilin did not change in response to poly(I:C) treatment in the cells expressing NLS-RL-CM (Figure S3C). We interpret these results to argue that nuclear RNA is generally required to maintain the integrity of Cajal bodies.

### Protein components of nuclear RNA granules do not co-assemble when nuclear RNA is degraded

We observed that the assemblies of the protein components of Cajal bodies, nuclear speckles and the nucleolus that formed when nuclear RNA was degraded were located in regions of the nucleus that were depleted in chromatin (Figure 4C and Figure S3). This observation suggested that the proteins of these different RNA granules could assemble or possibly aggregate together in the absence of their resident RNAs. We performed immunofluorescence analysis to determine if the protein components of the different nuclear RNA granules colocalized when nuclear RNA degradation was induced.

In cells in which nuclear RNA had been degraded, assemblies of coilin protein did not overlap with RPF1 foci but tended to be in close proximity with them (Figure 4D). The distinct nature of these new assemblies is consistent with those proteins, either individually or in combinations with other proteins, maintaining sufficient information to allow the formation of specific condensates. The coilin assemblies may be similar to structures referred to as nucleolar caps that contain coilin and form next to nucleoli when transcription is inhibited (Shav-Tal et al., 2005).

We also observed that the assemblies containing the nuclear speckle protein SRRM2 did not co-localize with the foci containing the Cajal body protein coilin or the nucleolar protein RPF1 (Figure 4D). Thus, these SRRM2 assemblies represent another distinct subassembly. However, the SRRM2 assemblies did overlap with a second nuclear speckle protein, SON.

Taken together, the ability of RNP granules to form novel assemblies in response to RNA degradation is consistent with these proteins having self-assembly properties either individually or with other proteins. In all the cases where we have observed de novo assemblies after RNA degradation, the proteins forming self-assemblies are thought to play roles in the assembly of their RNP granules (Hebert and Matera, 2000; Ilik et al., 2020). The observation that these new RNase-induced assemblies have different appearances, often being more spherical is consistent with the hypothesis that RNA can limit the self-assembly of some RNA binding proteins (Maharana et al., 2018).

In principle, RNA could be limiting the assembly of other RNA binding proteins to prevent the formation of inappropriate assemblies. This was suggested by the observation that FUS, an RNA binding protein with IDRs, formed assemblies after RNase A was injected into nuclei (Maharana et al., 2018). To examine if FUS behaved in a similar way after RNase L degradation of nuclear RNA, we monitored the distribution of FUS in cells with the NLS-RL-WT protein after poly(I:C) treatment.

We observed that FUS distribution was largely unchanged in the majority of cells with efficient nuclear RNA degradation. Specifically, FUS was dispersed throughout the nucleoplasm in the mock treated A549 cells (Figure 5A) and in 70% of cells responding to poly(I:C) treatment, we observed robust nuclear RNA degradation and FUS remained dispersed in the nucleoplasm (Figure 5A and 5B). This observation indicates that reduction of nuclear RNA does not lead to FUS self-assembly, at least at endogenous expression levels.

**Figure 5.**
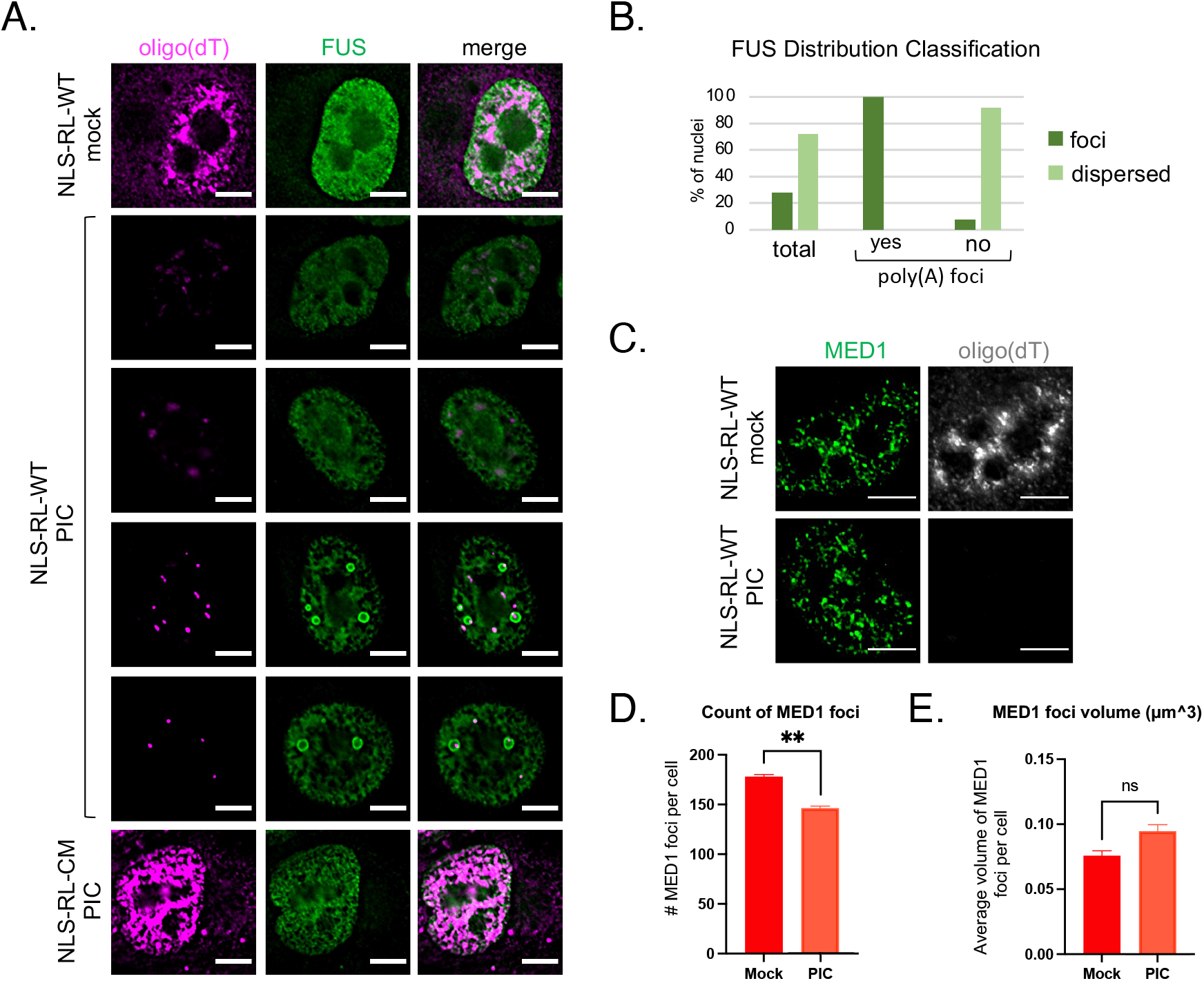
Analysis of the effect of nuclear RNA degradation on super-enhancer condensates and FUS assembly. Analysis of A549 cells with NLS-RL-WT or NLS-RL-CM either mock transfected or treated with poly(I:C) (PIC) for 5 hours. Scale bar 5 microns. A. FISH analysis to detect poly(A)+ RNA and IF analysis against FUS protein. Images are a single 0.2micron Z section. B. Graph of the percentage of nuclei with FUS dispersed in the nucleoplasm or in foci. All the nuclei (total) in which nuclear RNA was degraded (N=64) or the same nuclei classified as to whether or not they contained residual poly(A) foci. C. FISH analysis to detect poly(A)+ RNA and IF analysis against mediator complex protein, MED1, to detect super-enhancer condensates. D. Number of MED1 foci in mock and PIC treated NLS-RL-WT cells. E. Average volume of MED1 foci in mock and PIC treated NLS-RL-WT cells. D-E. Mean and SEM of 2 independent experiments with at least 10 cells of each cell type and condition per experiment. Mann-Whitney test, ns non-significant, ** P value ≤0.01.

Interestingly, in 30% of the responding cells, residual poly(A) foci appeared to nucleate FUS assemblies since FUS assemblies either overlapped with the poly(A) foci or assembled on their surface (Figure 5A and 5B). The assembly of the FUS foci was dependent on nuclear RNA degradation given that they were not observed after poly(I:C) treatment in cells with the catalytic mutant form of RNase L in the nucleus (Figure 5A). The observation that the FUS assemblies that form in the nucleus when RNA is degraded contain some type of RNA is consistent with these FUS assemblies requiring at least a low level of RNA. This suggests a model where FUS and RNA can assemble a complex condensate under certain concentrations of RNA and protein, with RNA being a required component of that assembly.

### Nuclear RNA degradation does not dramatically alter super-enhancer condensates

Recently, transcription factors and coactivators of transcription have been found to form condensates in the nucleus at the sites of super-enhancers (Sabari et al., 2018, Boija et al. 2019). Super-enhancers are clusters of enhancers that drive the robust expression of genes by assembling a high density of transcriptional machinery. Transcription of super-enhancers produces high quantities of what are referred to as enhancer RNAs or eRNAs (Hah et al., 2015). eRNAs have been proposed to promote enhancer function by playing a role in the phase separation of transcriptional components (Arnold et al. 2020), while high levels of RNA from nascent transcription are proposed to inhibit the formation of super-enhancer condensates (Henninger et al., 2021). We therefore examined whether super-enhancer condensates are altered when nuclear RNA is degraded by monitoring MED1, a component of the coactivator Mediator complex.

We observed that transcriptional condensates showed little alterations when nuclear RNA was degraded. MED1 was found in many small foci in the nucleus in mock treated cells (Figure 5C). There was a small but significant decrease in the number of MED1 foci in NLS-RL-WT cells in which nuclear RNA had been degraded, as identified by depletion of the nuclear oligo(dT) signal (Figure 5C and D). Although there was a tendency for nuclear RNA degradation to lead to a small increase in the average volume of MED1 foci in cells (Figure 5E), which would be consistent with nascent RNA inhibiting the formation of transcriptional condensates (Henninger et al., 2021), this difference was not significant.

The limited effect of nuclear RNA degradation on MED1 foci could be due to continued transcription of short-lived eRNAs. To test this possibility, we treated cells with Actinomycin D to inhibit transcription while we induced nuclear RNA degradation. Transcription inhibition did not further decrease the number of MED1 foci (Figure S4). The observation that nuclear RNA degradation did not disrupt MED1 foci suggests that eRNAs are not required to maintain condensates of transcriptional machinery.

### The role of RNA in maintaining membrane-less organelles

A striking aspect of this work is that many membrane-less organelles in both the nucleus and cytosol require RNA for their proper morphology and structure. The key observation is that degradation of either cytosolic or nuclear RNAs by activation of RNase L leads to loss of resident RNAs in these organelles and alterations in their morphology as judged by the localization of protein components. Since RNA affects the assembly of SGs, P-bodies, Cajal bodies, nuclear speckles, and the nucleolus we suggest that RNA molecules play a broad role in determining membrane-less compartments in eukaryotic cells.

In principle, RNA molecules can promote membrane-less organelle assembly in three manners (Figure 6). First, RNA molecules could allow for inter-molecular RNA-RNA interactions as suggested for the assembly of SGs (Van Treeck et al., 2018; Tauber et al., 2020). Alternatively, RNA molecules could provide scaffolding to increase the number and valency of proteins able to engage in intermolecular interactions. Finally, RNA molecules could serve as allosteric co-factors to alter the assembly properties of proteins that could drive organelle assembly. We anticipate that evolution will have made use of all these mechanisms in different contexts.

**Figure 6.**
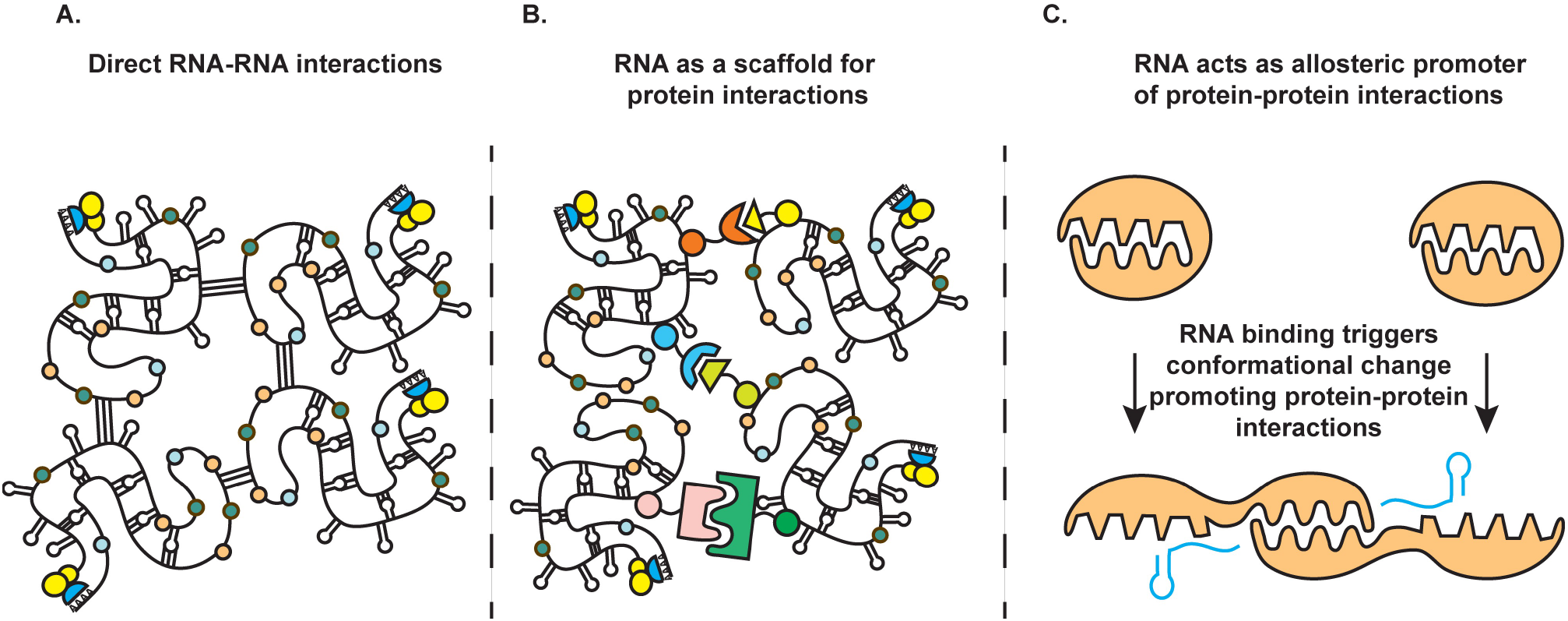
Model for the potential roles of RNA in the assembly and maintenance of membrane-less organelles containing RNA and protein.

For the RNA granules that are sensitive to the loss of RNA, we observe two different outcomes of their protein components with different implications. First, the proteins can distribute widely in their cellular compartment. For example, NPM1 and in many nuclei, RPF1, become widely distributed in the nucleoplasm when nucleolar RNAs are degraded (Figure 4). This dispersion of the marker protein argues that specific protein is insufficient to form any type of membrane-less assembly in the absence of RNA, and therefore can be inferred to have limited protein-based assembly parameters.

In other cases, we observed degradation of resident RNAs in a membrane-less organelle led to the formation of a new type of assembly. For example, when SGs disassemble after cytoplasmic RNA degradation, a residual membrane-less organelle, an RLB, remains with a distinctly different protein composition (Burke et al., 2020). Alternatively, in cells with complete nuclear RNA degradation, as assessed by oligo(dT) staining in the nucleus, coilin forms either a few large or multiple small irregular shaped assemblies (Figure 4). Similarly, when nuclear RNA is degraded, SRRM2 forms several large spherical assemblies (Figure 4). Although we cannot rule out the possibility that these new assemblies are being nucleated by residual RNA of some kind, it is likely that they reveal self-assembly properties of these proteins by themselves, or in conjunction with other nuclear proteins. Interactions with resident RNA molecules within granules or with RNA outside of their respective granules could limit proteins from associating with each other and in the absence of RNA, such protein-based assemblies can then accumulate. This demonstrates a second general rule of RNA molecules in modulating the organization of the cell which is to limit assembly of protein driven large assemblies that might be unproductive for normal biological function.

## MATERIALS AND METHODS

### Antibodies

For IF analysis: rabbit anti-Dcp1(D2P9W) antibody (CellSignaling 13233S) 1:500, mouse anti-DYKDDDDK tag(9A3) (Flag) antibody (Cell Signaling 8146S) 1:1000, mouse anti-G3BP antibody (AbCam ab56574) 1:1000, mouse anti-PDI antibody (CellSignaling 45596S) 1:500, rabbit anti-ZFP36L1/2 (TIS11B) antibody (Proteintech 12306-1-AP) 1:500, mouse anti-nucleophosmin (NPM1) (abcam ab10530) 1:1000, rabbit anti-RPF1 (Sigma HPA024642) 1:100, mouse anti-sc35 antibody (NovusBio NB100-1774) 1:1000, mouse anti-coilin antibody (abcam ab87913) 1:1000, rabbit anti-SRRM2 (ThermoFisher/Invitrogen PA5-66827) 1:200, rabbit anti-SON (ThermoFisher/Invitrogen PA5-65108) 1:200, rabbit anti-MED1 (abcam ab64965) 1:500, mouse anti-FUS antibody (atlas antibodies AMAb90549) 1:200, goat anti-rabbit IgG AF647 (abcam ab150079) 1:1000 and goat anti-mouse IgG FITC (abcam ab6785) 1:1000.

For Western analysis: mouse anti-RNase L antibody (Novus Biologicals NB 100-351) 1:2000, mouse anti-GAPDH-HRP (Santa Cruz Biotechnology sc47724) 1:5000, rabbit anti-Histone H3 antibody Novus Biologicals NB 500-171) 1:1000, anti-mouse IgG HRP (Sigma 4416) 1:5000, anti-rabbit IgG HRP (Cell Signaling 7074S) 1:5000.

### Cell culture and transfections

A549 and A549 RL-KO cell lines were described in Burke et al. 2019. Cells were maintained at 5% CO_2_ and 37°C in Dulbecco’s modified eagle medium (DMEM) supplemented with fetal bovine serum (10% v/v) and penicillin/streptomycin (1% v/v). Cells were tested for mycoplasma contamination by the cell culture core facility during the study and were negative.

### Oligonucleotides

cmyc_NLS_RL_sen primer 5’-CACCGGGACTGAAACTCGAGGGTACCGCCACCATGCCTGCTGCTAAGAGAGTGAAACTGGATGAGAGCAGGGATCATAACAA-3’

cmyc_NLS_RL anti primer 5’-ACAAGAAAGCTGGGTCTAGATTAGCACCCAGGGCTGGCCAACC-3’

oCDRP498 5’-CCATGACGGTGATTATAAAGATCATGACATCGACTACAAGGATGACGATGACAAGATGGAGGCGCTGAGTCG-3’

oCDRP499 5’-TAGGTTGTGGTTGTCTTTGTTCTTG-3’

oCDRP500 5’-ACAAAGACAACCACAACCTAATGGAGAGCAGGGATCATAACAAC-3’ oCDRP501 5’-ACAAGAAAGCTGGGTCTAGATCAGCACCCAGGGCTGGC-3’

### Plasmids

The cmyc NLS was fused to the N-terminus of RNase L and RNaseL-R667A by PCR amplification using primers cmyc_NLS_RL_sen and cmyc_NLS_RL anti and the RNase L and RNaseL-R667A pLenti-EF1-Blast plasmids described in Burke et al. 2019 as templates. The cmyc-NLS-RNaseL and RNaseL-R667A amplicons were inserted into Xho1/Xba1 cleaved pLenti-EF1-Blast vector using In-Fusion (Clontech). pVSV-G, pRSVRev and pMDlg-pRRE plasmids used for generation of lentiviral particles were described in Burke et al. 2019. Flag-tagged Dcp1a was fused to the RNase L by amplifying DCP1a using primers oCDRP498 and 499 and pcDNA3-FLAG-hDcp1a (Lykke-Andersen 2002) as a template and amplifying RNaseL and RNaseL-R667A using primers oCDRP500 and 501 and the RNase L and RNaseL-R667A pLenti-EF1-Blast plasmids described in Burke et al. 2019 as templates. The DCP1 and RNaseL amplicons were inserted into Xho1/Xba1 cleaved pLenti-EF1-Blast vector using In-Fusion (Clontech). All insert sequences were confirmed by sequencing.

### Generation of lentiviral particles and stable cell lines

To generate the NLS-RNase L, NLS-RNase L-R667A, DCP1-RNase L and DCP1-RNase L-R667A lentiviral particles, HEK293T cells (T25 flasks) were co-transfected with 2.7μg pLenti-EF1-cmycNLS-RNase L-Blast, pLenti-EF1-cmycNLS-RNase L-R667A-Blast, pLenti-EF1-DCP1-RNase L-Blast or pLenti-EF1-DCP1-RNase L-R667A-Blast, 0.8μg pVSV-G, 0.8μg pRSVRev, and 1.4μg pMDLg-pRRE using lipofectamine 2000 (ThermoFisher). Medium was replaced 6 hours post-transfection. Medium was collected at forty-eight hours post-transfection and filter-sterilized with a 0.45-um filter. To create NLS-RL-WT and NLS-RL-CM A549 stable cell lines, A549 cells were seeded in T-25 flasks, when 80% confluent, cells were incubated for 1 hour with 1 mL of either NLS-RNase L or NLS-RNase L-R667A lentiviral particles containing 10 μg of polybrene with periodic rocking. Normal medium was then added to the flask and incubated for twenty-four hours. Medium was removed 24 hours post-transduction and replaced with selective growth medium containing 5 μg /ml of Blasticidine S hydrochloride (Sigma-Aldrich). Selective medium was changed every few days then replaced with normal growth medium after 7 days. DCP1-RNase L and DCP1-RNase L-R667A A549 RL-KO stable cells lines were made in a similar manner.

### Cell fractionation and western blotting

Fractionation of A549, A549-RL-KO, A549 NLS-RL-WT and A549 NLS-RL-CM cells was performed as described in Burke and Sullivan 2017. Cells grown in one well of 6 well plate were trypsinized, equivalent number of cells were washed with phosphate-buffered solution (PBS), resuspended in 200μl of CSKT buffer [10mM PIPES (pH 6.8), 100mM NaCl, 300mM sucrose, 3mM MgCl_2_, 1mM EDTA, 1mM DTT, 0.5% (vol/vol) TritonX-100, protease inhibitors (Roche)], 100 μl was removed for whole cell lysate and mixed with 100 μl SDS lysis buffer (1%SDS, 2% 2-mercaptoethanol), the remaining 100μl was placed on ice for 10 minutes. Nuclei were then pelleted by centrifugation at 5000×g for 5 minutes. The supernatant 100 μl cytosolic fraction was removed and mixed with 100μl SDS lysis buffer. The nuclei were then washed in 100 μl CSKT buffer and centrifuged at 5000×g for 5 minutes. The nuclei were then resuspended in 100 μl CSKT buffer followed by addition of 100μl SDS lysis buffer to make the nuclear fraction. Samples were boiled for 10 minutes followed by vortexing for 30 seconds. Equal volumes of each fraction and whole cell lysate were separated on 4-12% NuPAGE gel (ThermoFisher) with PAGEruler prestained protein markers (Thermofisher), transferred to Protran membrane (Amersham), the blot was cut into 3 pieces based on prestained protein markers. The strip with proteins 70kD and larger was incubated with mouse anti-RNase L antibody mouse. The strip with proteins 70kD to 25kD was incubated with anti-GAPDH-HRP. The strip with proteins 25kD and smaller was incubated with anti-Histone H3 antibody. After incubation with appropriate HRP conjugated secondary antibodies the blots were developed using SuperSignal West Dura substrate (Pierce). Images of blots obtained with ImageQuant LAS 4000 (GE Healthcare). Western blotting using anti-RNase L antibody against DCP1-RNase L fusion proteins was performed on whole cell lysates prepared as described above.

### Sequential immunofluorescence and FISH analysis following transfection with poly(I:C)

Cells were seeded on sterile coverslips in 6 well plates, grown to 40-60% confluence, then transfected with poly(I:C) HMW (InvivoGen: tlrl-pic) using 3μl of lipofectamine 2000 (Thermo Fisher Scientific) per 1μg of poly(I:C) or mock transfected using lipofectamine alone for 4 to 5 hours. Actinomycin D, when used, was added to final concentration of 1μg/ml at the same time as poly(I:C) addition. Sequential immunofluorescence and FISH analysis was performed following manufacturer’s protocol (https://biosearchassets.blob.core.windows.net/assets/bti_custom_stellaris_immunofluorescence_seq_protocol.pdf) with the following changes, 4% paraformaldehyde in 1XPBS was used for fixation, antibody incubations were performed in 0.1ml 1% nuclease free BSA (Invitrogen) 1XPBS after blocking for 1 hour in same solution, Dapi staining was performed in PBS following Wash Buffer B step, and coverslips were mounted using Prolong Glass (Invitrogen). FISH probes: human GAPDH with Quasar 670 dye (Stellaris SMF-2019-1), human MALAT1 with Quasar 670 dye (Stellaris SMF-2046-1), human snoRD3A probe GCGTTCTCTCCCTCTCACTCCCCAATA-AlexaFluor647 and oligod(T)30-Cy3 were purchased from IDT.

### Microscopy and image analysis

Immunofluorescence and FISH with Dapi staining were imaged using a wide-field DeltaVision Elite microscope with an Olympus UPlanSApo 100×/1.40-NA oil objective lens and a PCO Edge sCMOS camera with appropriate filters at room temperature using SoftWoRx Imaging software taking 15 Z planes at 0.2μm/section per image. Deconvolved images were processed using ImageJ with FIJI package. Max projections of Z-planes were performed, and brightness and contrast adjusted for each channel to best view results. Super resolution images of TIS granules were obtained using a Nikon CSU W1 SoRa microscope with 100X/1.45 NA oil objective lens and Photometrics Prime BSI sCMOS Camera. Quantification of SGs, PBs, TIS granules and MED1 foci was performed using Imaris Image Analysis Software (Bitplane) (University of Colorado-Boulder, BioFrontiers Advanced Light Microscopy Core) using non-deconvolved images.

## Abbreviations

DFC: dense fibrillar component
eRNA: enhancer RNA
FC: fibrillar center
GC: granular component
lncRNA: long non-coding RNA
NPM1: nucleophosmin 1
PBs: processing bodies
PDI: protein disulfide isomerase
RPF1: ribosome production factor 1
SGs: stress granules
ActD: Actinomycin D

## AUTHOR CONTRIBUTIONS

C.J.D, J.M.B and R.P conceptualization, C.J.D data curation, C.J.D, P.K.M and J.M.B. investigation, C.J.D. formal analysis. C.J.D. visualization, C.J.D and R.P. writing-original draft. C.J.D, J.M.B, P.K.M and R.P writing-review and editing, R.P. supervision, J.M.B. and R.P. funding acquisition.

## ACKNOWLEDGEMENTS

We thank Evan Lester for sharing reagents. We thank Theresa Nahreini, Facility Manager, Cell Culture Facility, Biochemistry for her expertise and support. The Analysis Workstation and the Imaris software package at the BioFrontiers Institute Advanced Light Microscopy Core were supported by NIH 1S10RR026680-01A1. This work was supported by the National Institute of Allergy And Infectious Diseases of the National Institutes of Health under Award Number F32AI145112 (J.M.B) and Howard Hughes Medical Institute (C.J.D. and R.P.). The authors declare no competing financial interests.

## Notes

### Competing Interest Statement

The authors have declared no competing interest.

