## Supplementary material for "RNA is required for the maintenance of multiple cytoplasmic and nuclear membrane-less organelles": Figure S1

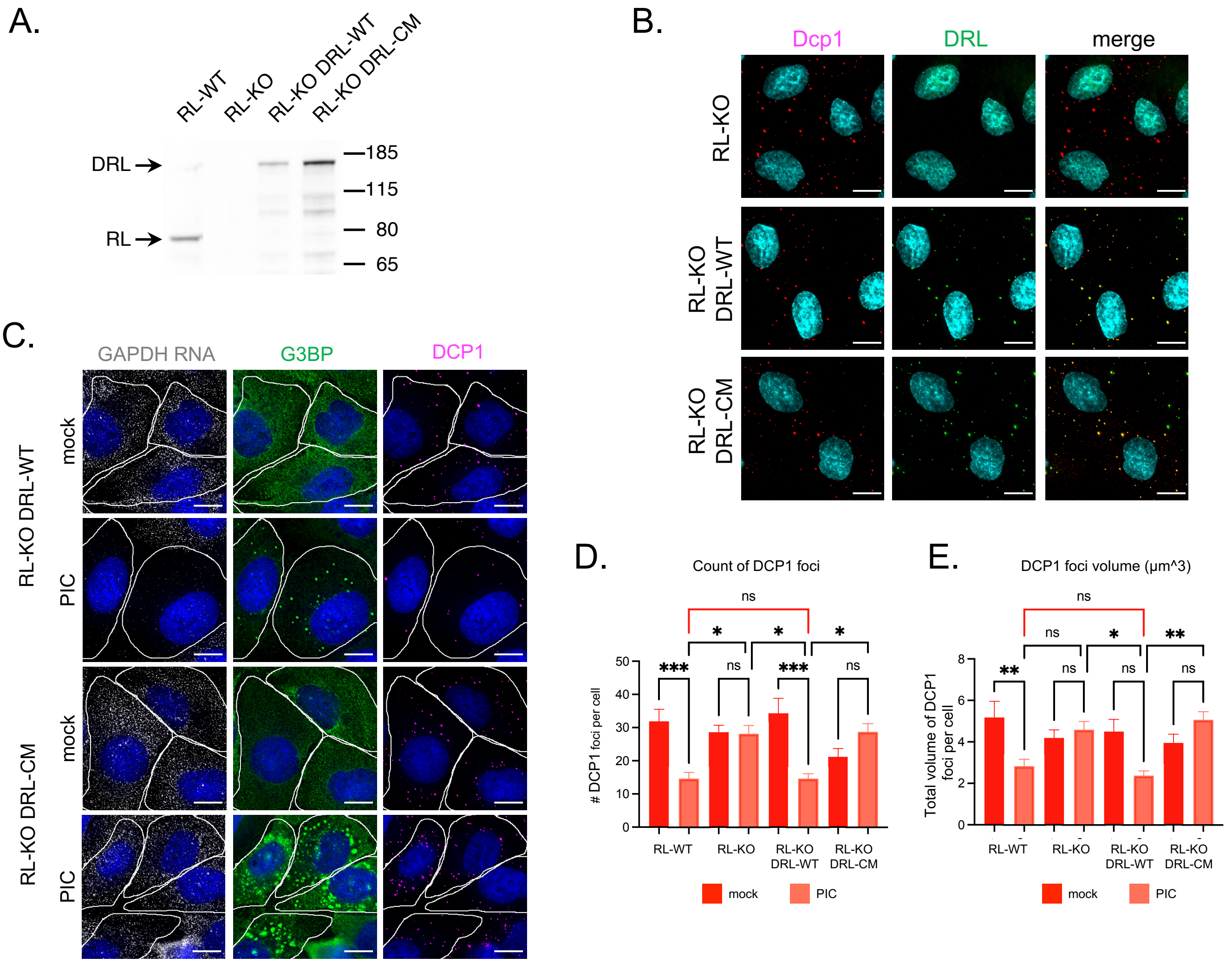


Figure S1. Targeting RNase L to P-bodies did not increase the effect of RNase L activation on P-body number or total volume. A. Full-length Dcp1 RNase L (DRL) fusion proteins are expressed. Western analysis using anti-RNase L antibody of whole cell lysates from A549 (RL-WT) cells, A549 RNase L knock out cells (RL-KO) and RL-KO cells transduced with lentiviral vectors containing either wild-type RNase L fused to DCP1a (RL-KO DRL-WT) or catalytic mutant RNase L-R667A fused to DCP1a (RL-KO DRL-CM). Arrows on left indicate migration of the endogenous RNase L (RL) and the DCP1 RNase L fusion proteins (DRL). B. DRL fusion proteins co-localize with P-bodies. IF analysis using anti-DCP1b antibody to detect P-bodies (DCP1) or anti-Flag antibody to detect Flag-tagged DCP1a-RNase L fusion proteins. Scale bar 10 microns. C. DRL-WT fusion protein is active. GAPDH smFISH and IF analysis using anti-G3BP antibody (G3BP) and anti-Dcp1b antibody (DCP1) in A549 RL-KO cells with DRL-WT or DRL-CM fusion proteins either mock transfected (mock) or transfected with poly(I:C) (PIC) for 5 hours. Scale bar 10 microns. D. Number of DCP1 foci in mock and PIC treated cells (mean and SEM). E. Total volume of DCP1 foci in mock and PIC treated cells (mean and SEM). D-E. At least 10 cells were counted per cell type per condition. One way ANOVA with Tukey's multiple comparison, ns non-significant, * P value $\leq$0.05, ** P value $\leq$0.01, *** P value $\leq$0.0001. Note: no significant difference in # of DCP1 foci or total volume of DCP1 foci between RL-WT and RL-KO DRL-WT after PIC treatment.
