## Supplementary material for "RNA is required for the maintenance of multiple cytoplasmic and nuclear membrane-less organelles": Figure S2

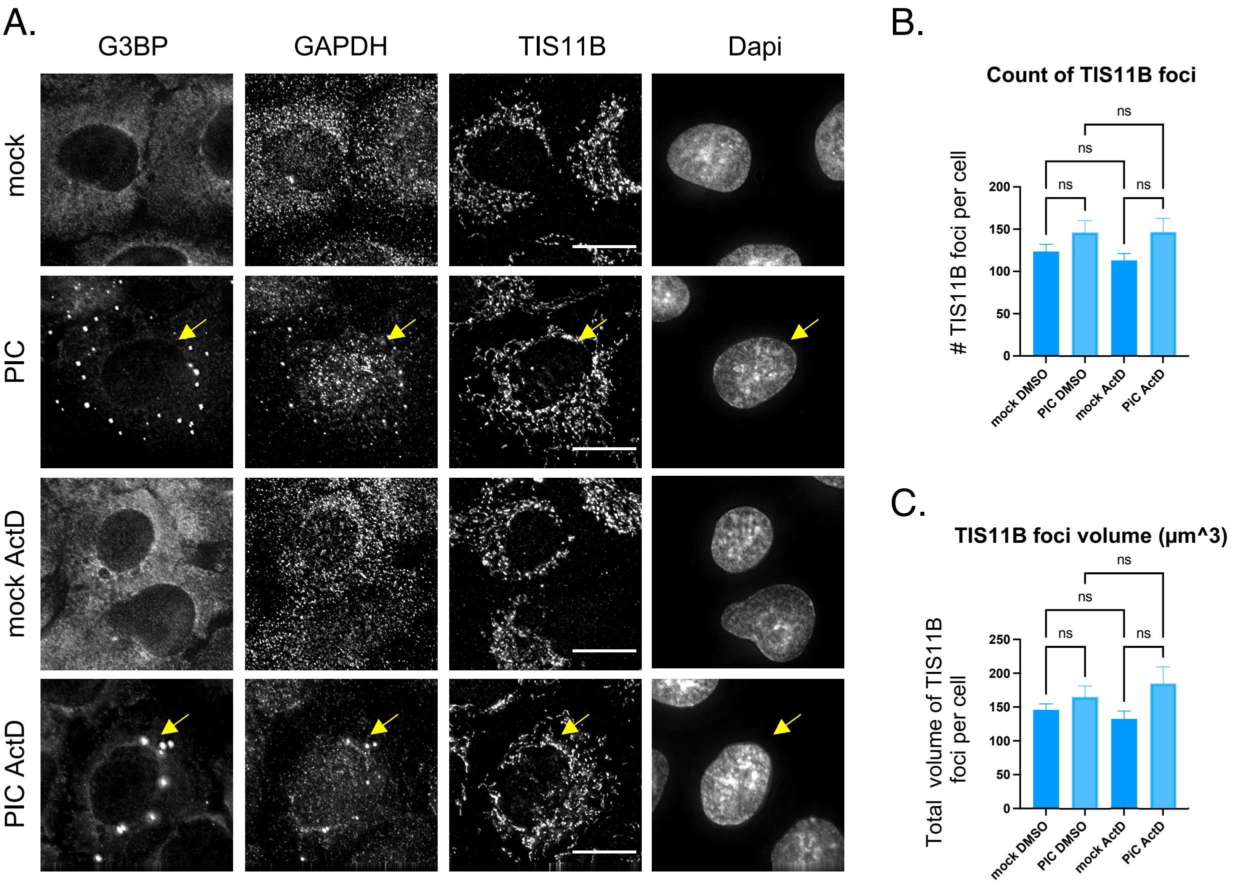


Figure S2. Inhibition of transcription in combination with activation of RNase L in the cytoplasm did not alter TIS granules.

A. IF analysis of TIS11B in A549 cells mock transfected or treated with poly(I:C) with DMSO or 1µg/ml Actinomycin D (ActD) for 5 hours. IF against G3BP used to monitor cells responding to poly(I:C), GAPDH FISH analysis used to monitor cytoplasmic RNA degradation and transcription inhibition. Yellow arrows indicate cells responding to poly(I:C). Scale bar 5 microns. B. Number of TIS11B foci per cell in mock, PIC, ActD or PIC and ActD treated cells (mean and SEM). C. Average volume of TIS11B foci per cell in mock, PIC, ActD or PIC and ActD treated cells (mean and SEM). B-C. At least 10 cells in each condition were counted. One way ANOVA with Tukey's multiple comparison, ns non-significant.
