## Supplementary material for "RNA is required for the maintenance of multiple cytoplasmic and nuclear membrane-less organelles": Figure S3

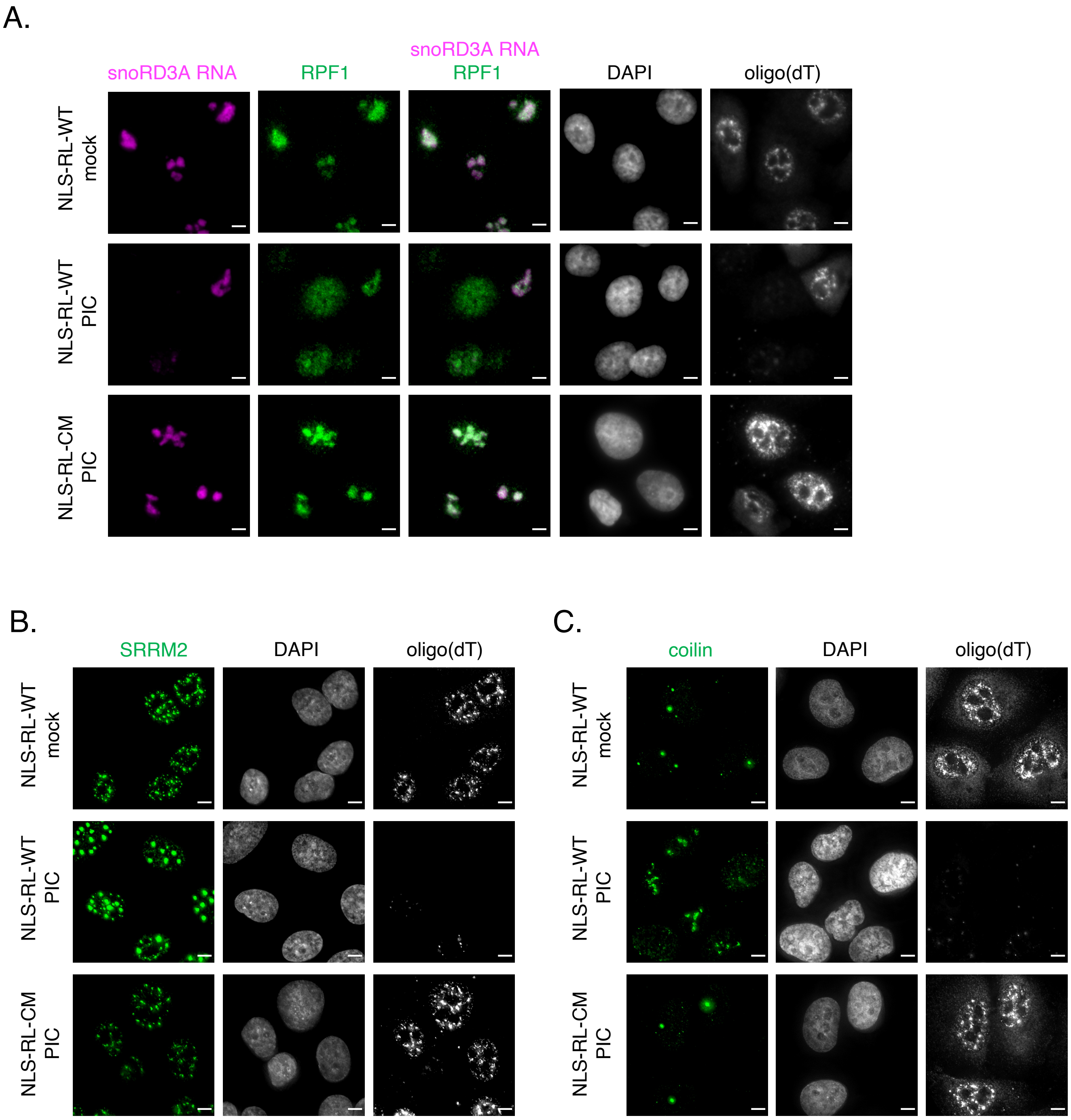


Figure S3 . Changes in nuclear RNA granule morphology is dependent on loss of nuclear RNA.
IF analysis of nuclear RNA granule proteins in A549 cells expressing nuclear-localized wildtype RNase L or catalytic mutant RNase L-R667A mock transfected or treated with poly(I:C) for 5 hours. Scale bar 5 microns. A. FISH analysis to detect nucleolar-localized snoRD3A RNA and poly(A)+ RNA. IF analysis of nucleolar protein RPF1. B. Oligo(dT) FISH and IF analysis of nuclear speckle protein SRRM2 with sc35 antibody. C. Oligo(dT) FISH and IF analysis of Cajal body protein coilin.
