## Supplementary material for "RNA is required for the maintenance of multiple cytoplasmic and nuclear membrane-less organelles": Figure S4

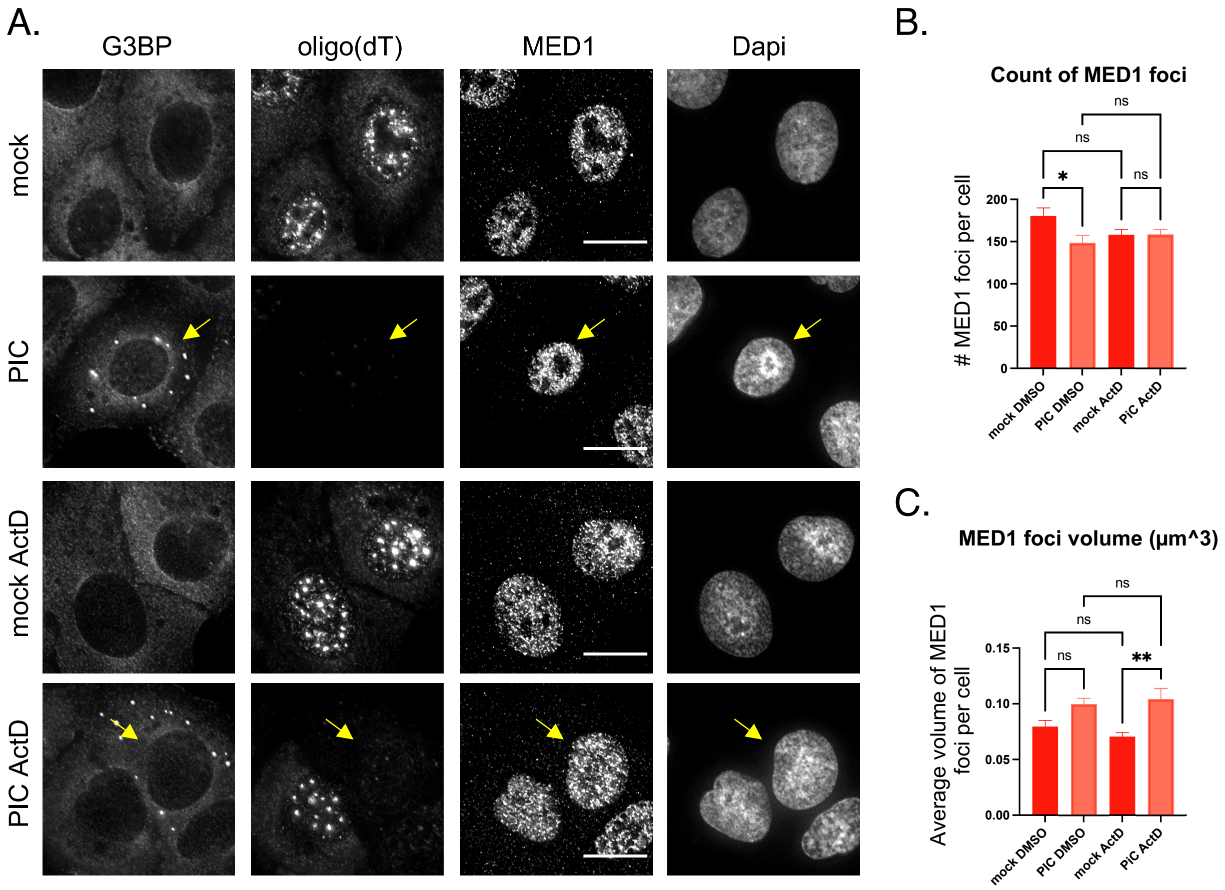


Figure S4. Inhibition of transcription did not enhance the decrease in the number of MED1 foci that occurred in response to nuclear RNA degradation.
A. IF analysis of super enhancer condensate protein MED1 in A549 cells expressing nuclear-localized wildtype RNase L mock transfected or treated with poly(I:C) with DMSO or 1μg/ml Actinomycin D (ActD) for 5 hours. IF against G3BP used to monitor cells responding to poly(I:C), FISH analysis for poly(A)+ RNA used to monitor nuclear RNA degradation and transcription inhibition. Yellow arrows indicate cells responding to poly(I:C). Scale bar 5 microns. B. Number of MED1 foci per cell in mock, PIC, ActD or PIC and ActD treated cells (mean and SEM). C. Average volume of MED1 foci per cell in mock, PIC, ActD or PIC and ActD treated cells (mean and SEM). B-C. At least 10 cells in each condition were counted. One way ANOVA with Tukey's multiple comparison, ns non-significant, * P value $\leq 0.05$, ** P value $\leq 0.01$.
