## Supplementary material for "RNA is required for the maintenance of multiple cytoplasmic and nuclear membrane-less organelles": Table S1

**Table S1. Copy number and change in RNA levels after RNase L activation for RNAs reported to co-localize with TIS granules[a].**

| mRNA name | molecules/cell[b] | RL-WT PIC vs mock log2 fold change[c] |
|---|---|---|
| CD47 | 7 | -1.61 |
| BCL2 | 2 | 0.08 |
| CD274 | 5 | 1.66 |
| ELAV1 | 25 | -1.35 |
| CCND1 | 139 | -0.89 |
| FUS | 35 | -0.67 |

[a]Ma, W., and Mayr, C. 2018. A membraneless organelle associated with the endoplasmic reticulum enables 3'UTR-mediated protein-protein interactions. Cell 175:1492-1506.

[b]Khong, A., Matheny, T., Jain, S., Mitchell, S.F., Wheeler, J.R., and Parker, R. 2017. The stress granule transcriptome reveals principles of mRNA accumulation in stress granules. Mol. Cell 68:808-820.e5.

[c]Burke, J.M., Moon, S.L., Matheny, T., and Parker, R. 2019. RNase L reprograms translation by widespread mRNA turnover escaped by antiviral mRNAs. Mol. Cell 75:1203-1217.
